# HSP90 inhibition modulates NFκB signaling in airway goblet cell metaplasia

**DOI:** 10.1101/2020.05.24.113902

**Authors:** Rosarie A. Tudas, Ryan M. Gannon, Andrew L. Thurman, Mallory R. Stroik, Joseph Zabner, Alejandro A. Pezzulo

## Abstract

Goblet cell metaplasia and mucus hyper-production are key features of chronic muco-obstructive lung diseases such as asthma, chronic bronchitis, and cystic fibrosis. Various mechanisms lead to goblet cell metaplasia in the airways; the driving mechanism for goblet cell metaplasia in a specific patient may be unknown. We recently found that heat shock protein 90 (HSP90) is important for both IL-13- and IL-17- induced airway goblet cell metaplasia. HSP90 interacts with multiple clients that are important in goblet cell metaplasia including Akt, Jak/STAT, IRS, Notch, and various kinases involved in NFκB signaling. Here, we used a targeted phospho-proteomic approach to identify candidate HSP90 clients modulated by the HSP90-inhibitor geldanamycin. NFκB family members were enriched amongst the top candidate targets of HSP90 inhibition in IL-13 an organotypic model of human airway epithelia. We hypothesized that HSP90 inhibition modulated goblet cell metaplasia by interfering with NFκB signaling. We used transcription factor activation, nuclear translocation, and phospho-specific immunofluorescence assays to investigate how IL-13 exposure and HSP90 inhibition modulated NFκB. We found that HSP90 inhibition prevented goblet cell metaplasia by non-canonically blocking NFκB p100/p52 function in human airway epithelia. NFκB modulation via its interaction with HSP90 is a pharmaceutically feasible therapeutic target for goblet cell metaplasia; this approach may enable treatment of patients with chronic muco-inflammatory lung diseases with both known or unidentified disease-driving mechanisms.

## Introduction

Chronic muco-obstructive lung diseases such as asthma, chronic bronchitis, and cystic fibrosis are a major cause of mortality and disability worldwide (1–3). There are currently no curative treatments for most patients with chronic muco-obstructive diseases.

While the mechanisms and triggers of different chronic muco-obstructive lung diseases are varied, they share as a hallmark the accumulation of mucus-producing goblet cells (goblet cell metaplasia) (4). Goblet cell metaplasia is associated with mucus hyper-production and hyper-secretion, and disproportionally contributes to the disabling effects of chronic lung diseases. Treatments to modulate goblet cell metaplasia are promising approaches to improving the lives of people with muco-obstructive lung diseases.

Various environmental triggers can activate pathways that cause airway goblet cell metaplasia (5–8). One pathway that has been well characterized is the T helper 2 (T_h_2) pathway in which interleukin 13 (IL-13) activates downstream signaling that ultimately results in goblet cell metaplasia. T_h_2 signaling drives disease in approximately half of people with asthma (9–12). T_h_17 signaling drives goblet cell metaplasia in some people with asthma, chronic bronchitis, and cystic fibrosis (10). In many patients with chronic airway goblet cell metaplasia, the driving mechanism is unknown.

Heat shock protein 90 (HSP90) is important for both IL-13- and IL-17-induced goblet cell metaplasia. We previously showed that HSP90 inhibition blocks and reverts IL-13- and IL-17-induced goblet cell metaplasia in human airway epithelia cells *in vitro* and in mice *in vivo* (13). HSP90 is a chaperone protein required in the proper folding and stabilization of hundreds of client proteins. HSP90 inhibitors act on the ATPase activity of HSP90 and results in ubiquitination and proteasomal degradation of its clients (14). Various HSP90 clients are important in goblet cell metaplasia, including Akt (15,16), Jak/STAT (17), IRS (18,19), Notch (20), and kinases important for NFκB signaling (21–25).

The key HSP90 client in goblet cell metaplasia targeted by HSP90 inhibition is unknown. Here, we used a targeted phospho-proteomic approach to identify candidate targets modulated by the HSP90 inhibitor geldanamycin implicated in IL-13-induced goblet cell metaplasia; NFκB family members were prominently enriched amongst the candidate targets. The NFκB family proteins are critical inflammatory signaling pathway in multiple immune and non-immune cell types and are regulated via multiple mechanisms (26–29). We directly examined the role of specific NFκB subunits in goblet cell metaplasia using DNA-binding, nuclear translocation, and immunofluorescence microscopy assays. We found that HSP90 inhibition affects NFκB signaling non-canonically to prevent goblet cell metaplasia in human airway epithelia in vitro.

## Results

### HSP90 inhibition modulates phosphorylation of several proteins in IL-13-treated human primary epithelia

We have previously shown that HSP90 inhibition blocks and partially reverts IL-13-induced goblet cell metaplasia in primary human airway epithelia *in vitro* (13). HSP90 is known to interact with many client proteins. To determine HSP90-interacting proteins essential in goblet cell metaplasia, we used an antibody array (Full Moon BioSystems Phospho-Explorer Array) consisting of 1318 site-specific and phospho-specific protein targets from over 30 signaling pathways. Primary airway epithelial cells from human donors were cultured at the air-liquid interface and exposed to 20 ng/ml of IL-13 for 28 days to establish goblet cell metaplasia. With continued IL-13 treatment, an HSP90 inhibitor (HSP90i, 25μM geldanamycin), was then added for 1, 2, 7, or 14 days. This dosage of IL-13 and geldanamycin was used in all experiments and was previously determined (13). To measure modulation of protein phosphorylation, we performed a Phospho-Explorer antibody array and evaluated the effect of HSP90 inhibition on the phosphorylation of numerous proteins in IL-13-exposed airway epithelial cells. The ratio of phosphorylated to total protein was calculated for each phosphorylation site included on the array. We found that phosphorylation of several proteins in IL-13-exposed airway epithelial cells were modulated by HSP90 inhibition (Fig. 1 and Supporting Table 1). We identified 46 proteins whose phosphorylation was up- or down-regulated by HSP90 inhibition by over two-fold (log_2_) relative to IL-13-only control. These data show that HSP90 inhibition affects multiple signaling pathways in IL-13-stimulated human airway epithelia.

**Figure 1.**
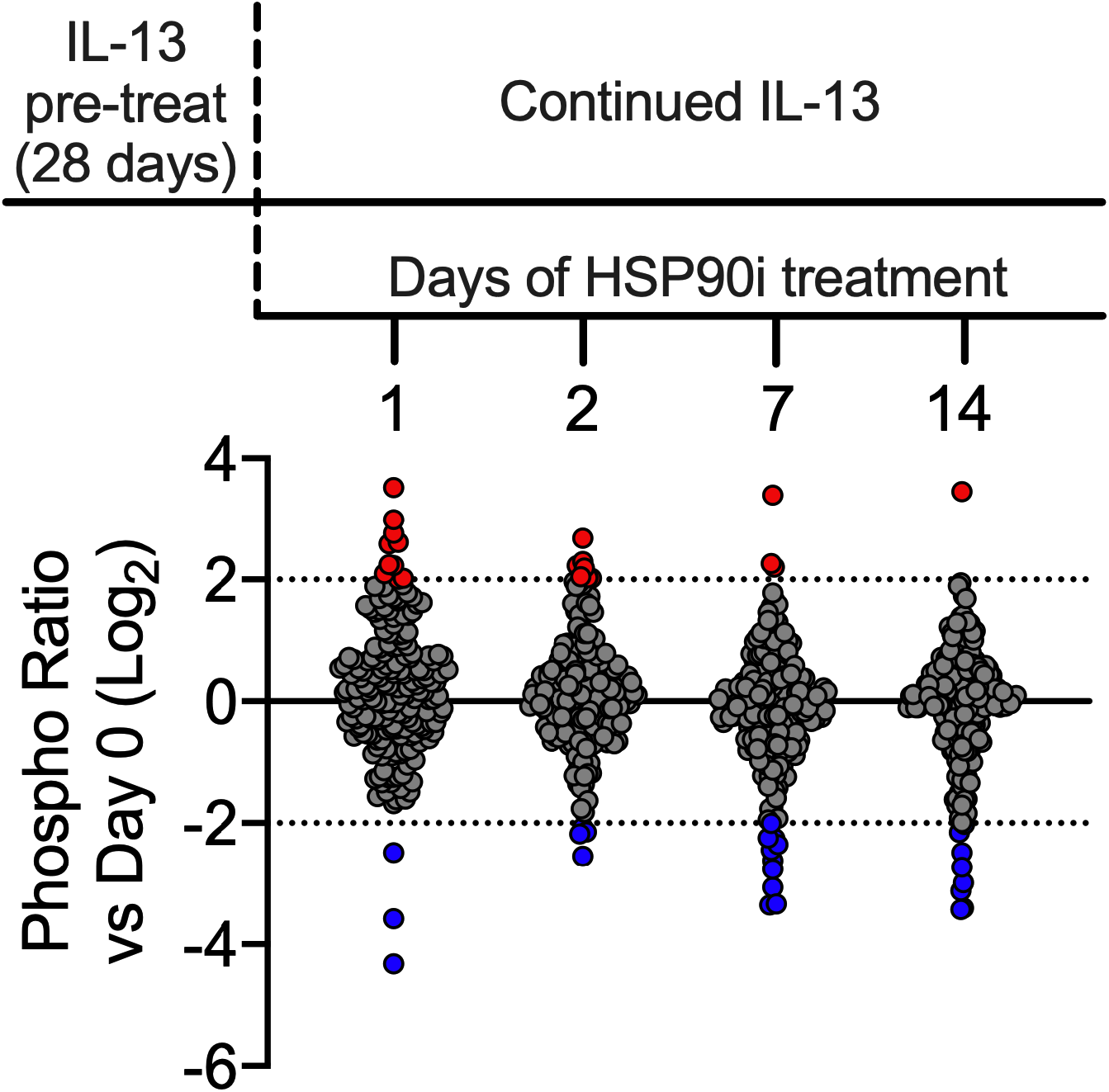
HSP90 inhibition modulates phosphorylation of several proteins in IL-13-treated human primary epithelia. Primary human airway epithelia *in vitro* were exposed to IL-13 (20ng/ml) for 28 days. An HSP90 inhibitor, geldanamycin (25μM), was then added for 1, 2, 7, or 14 days. A Phospho-Explorer antibody array was performed to determine modulation of protein phosphorylation by HSP90 inhibition in IL-13-treated human airway epithelial cells. Ratio of phosphorylated to total protein was obtained for each phosphorylation site. Data is shown in log_2_ fold change compared to IL-13-only treatment. Each dot represents a phosphorylation site on a protein. The top phosphorylated sites are shown in red, and the top dephosphorylated sites are shown in blue.

### Protein phosphorylation profiling identifies NFκB as a candidate HSP90 client modulating IL-13- induced goblet cell metaplasia

The HSP90 inhibitor-induced phosphorylation changes in IL-13-exposed airway epithelial cells were ranked by their effect magnitude. We identified the top ten modulated protein-phosphorylation sites (Fig. 2 and Supporting Table 1). While many proteins were affected by HSP90 inhibition, the top targets were enriched in NF*κ*B subunits. These data show that NF*κ*B phosphorylation is modulated by IL-13 in human airway epithelial cells, and suggest that NFκB is involved in L13-induced airway goblet cell metaplasia.

**Figure 2.**
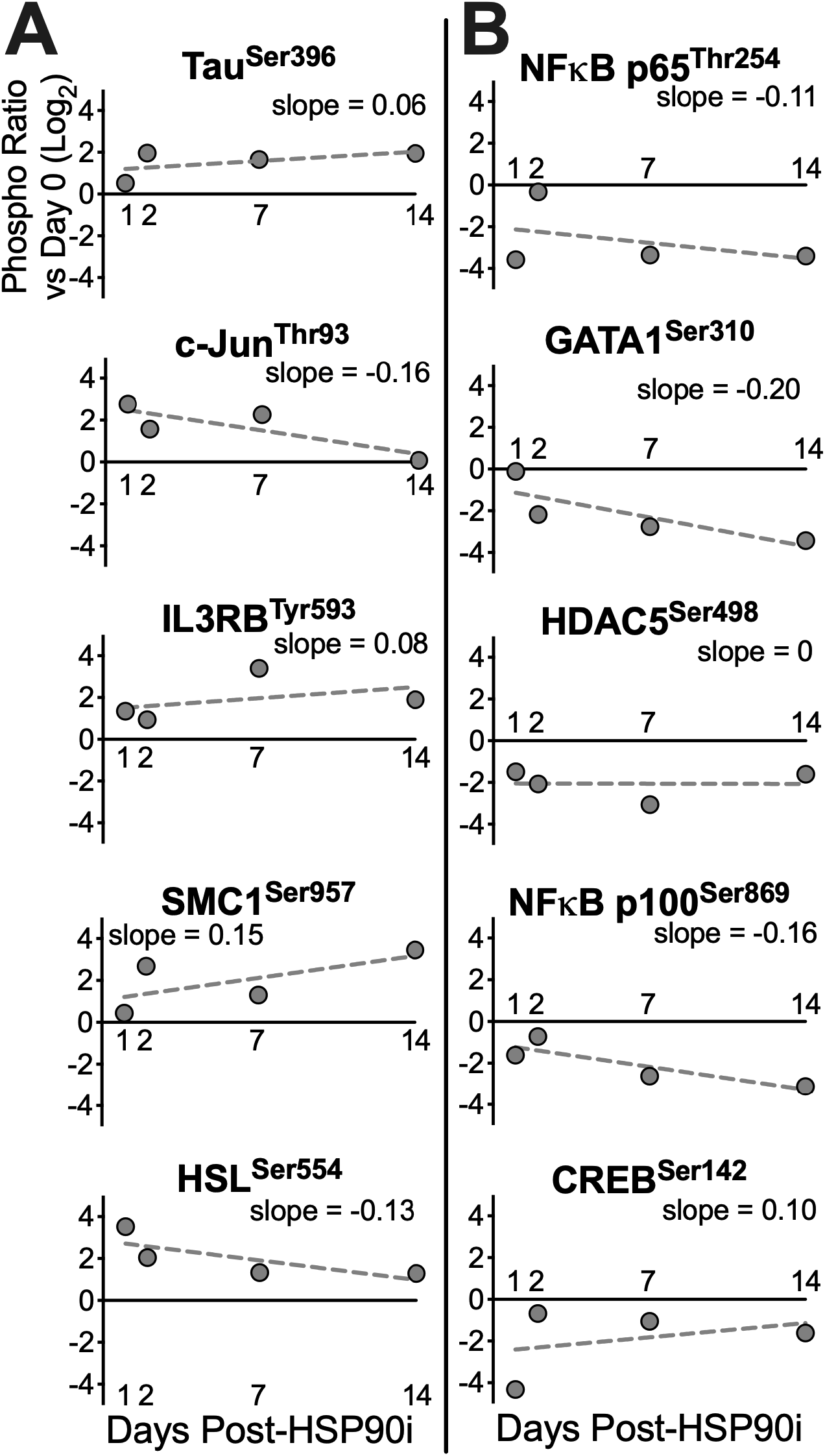
Protein phosphorylation profiling identifies NF*κ*B as a candidate HSP90 client modulating IL-13-induced goblet cell metaplasia. The top protein-phosphorylation sites modulated by HSP90 inhibition in IL-13-exposed airway epithelial cells, as measured in the antibody array, are listed. The top five phosphorylated protein-phosphorylation sites are shown in red, and the top five dephosphorylated protein-phosphorylated sites are shown in blue.

### IL-13 and HSP90 inhibition have both synergistic and antagonistic effects on phosphorylation of NFκB subunits

To determine whether HSP90 inhibition blocks IL-13-induced phosphorylation changes on NFκB subunits, we exposed primary human airway epithelial cells to vehicle, IL-13, HSP90 inhibitor, or IL-13 plus geldanamycin for 2 days. The same antibody array was used to evaluate phosphorylation of specific residues on NF*κ*B subunits p100/p52, p105/p50, and p65 (or RelA). The ratio of phosphorylated to total protein was calculated for each phosphorylation site of the NF*κ*B subunits included in the antibody array. The phospho-total ratios were then compared against baseline ratios.

The effects of geldanamycin and IL-13 on NF*κ*B were subunit- and phosphorylation site-dependent (Supporting Table 2). We observed several patterns of synergism and antagonism between geldanamycin and IL-13 effects on NF*κ*B phosphorylation sites. Some phosphorylation sites were predominantly affected by geldanamycin (Fig. 3A) or IL-13 (Fig. 3B) even when combined. Interestingly, geldanamycin and IL-13 acted antagonistically (Fig. 3C) or synergistically (Fig. 3D) on some phosphorylation sites. Lastly, IL-13, geldanamycin, and the combination of both also showed analogous effects on some NF*κ*B phosphorylation sites (Fig. 3E).

**Figure 3.**
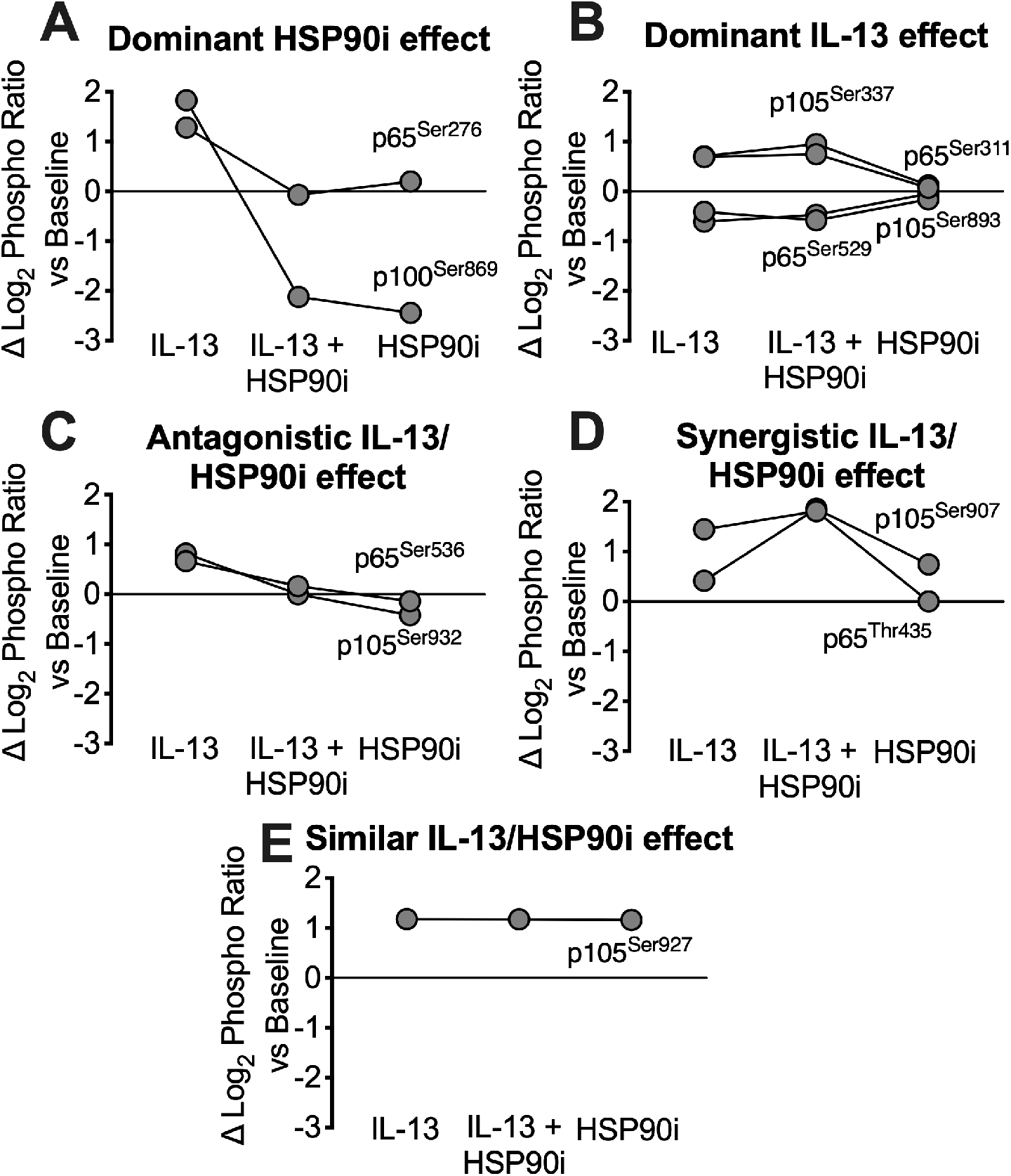
Patterns of effect of IL-13 and HSP90 inhibition in phosphorylation of various NF*κ*B subunits in primary human airway epithelia *in vitro*. Primary human airway epithelia *in vitro* were treated with vehicle, IL-13, HSP90 inhibitor, or IL-13 and HSP90 inhibitor for 2 days. HSP90 inhibition effects on IL-13-induced phosphorylation changes on NF*κ*B subunits were evaluated using a Phospho Explorer antibody microarray. Ratio of phosphorylated to total protein was obtained for each phosphorylation site. Data is shown in log_2_ fold change from baseline. Phosphorylation sites on NF*κ*B subunits are represented by a line which are organized into five patterns: (A) HSP90 inhibitor effect was dominant, (B) IL-13 effect was dominant, (C) IL-13 and HSP90 inhibitor have antagonistic effects, (D) IL-13 and HSP90 inhibitor have synergistic effect, and (E) IL-13 and HSP90 inhibitor have similar effects.

Phosphorylation of NF*κ*B is a regulatory mechanism that can have different downstream effects depending on the residues that are modified. Phosphorylation on some NF*κ*B residues may induce processing of the inactive form to its active form that can translocate into the nucleus and affect transcription, as in the case of p100/p52 Serine-869 (S869) (30). Our data show that IL-13 induces an increase in phosphorylated (p) S869 on p100/p52 subunit while HSP90 inhibition results in a decrease in p100/p52 pS869 compared to untreated control. These data suggest that IL-13 induces the activation of NF*κ*B in airway epithelial cells *in vitro* which may affect transcription of genes involved in airway goblet cell metaplasia. Moreover, HSP90 inhibition blocks this effect of IL-13 which is consistent with our previous data showing HSP90 inhibition blockade of IL-13-induced goblet cell metaplasia *in vitro.* NF*κ*B phosphorylation can also inhibit constitutive NFκB processing by stabilization of the inactive form thereby sequestering it in the cytosol, as in the case of p105/p50 Serine-907 (30). In our cultured airway epithelial cells, HSP90 inhibition increases p105/p50 pS907. Interestingly, IL-13 treatment results in a similar effect, and treatment with both IL-13 and geldanamycin appear to have a synergistic effect.

Phosphorylation at specific NF*κ*B residues may also be essential for DNA binding as has been shown for p105/p50 Serine-337 (31). Here, we observed that in airway epithelial cells *in vitro*, IL-13 increases p105/p50 pS337. However, HSP90 inhibition does not seem to block this effect. This inability of geldanamycin to block IL-13-induced up- or down-regulation of phosphorylated NF*κ*B is observed in several NF*κ*B residues. This further implicates the NF*κ*B pathway in IL-13-induced airway goblet cell metaplasia. Moreover, this also suggests that HSP90 may play a role in the phosphorylation of some, but not all, residues of the NF*κ*B subunits. Thus, HSP90 inhibition may be blocking and reverting IL-13- induced goblet cell metaplasia via modulation of specific NF*κ*B subunits.

### IL-13 does not induce DNA binding of p52 subunit in human airway epithelial cells in vitro

The observed IL-13- and HPS90i-effects on NF*κ*B phosphorylation in our antibody array data suggest that several NF*κ*B subunits may be involved in IL-13-induced goblet cell metaplasia. However, our data also suggests that HSP90 inhibition is acting via modulation of specific NF*κ*B residues. Since p100 pS869 was the most differentially modulated NF*κ*B subunit by IL-13 and geldanamycin, we examined the DNA binding capacity of its processed active form, p52. Based on our findings of IL-13-induced phosphorylation of p100 S869, we expected subsequent p100 processing into p52 after IL-13 exposure. Therefore, we hypothesized that IL-13 treatment will induce DNA binding of p52 in human airway epithelial cells. Since NF*κ*B activation has been shown to be transient due to tight regulation via negative feedback (32), exposure duration was reduced to 1 hour. Human airway epithelial cells *in vitro* were treated with vehicle or IL-13 for 1 hour, and nuclear extracts were collected. To measure p52 DNA binding, we used the TransAM DNA-binding ELISA. We found that IL-13 did not induce DNA binding of p52 (Fig. 4). This suggests that NFκB modulation by IL-13 does not induce canonical p100/p52 signaling.

**Figure 4.**
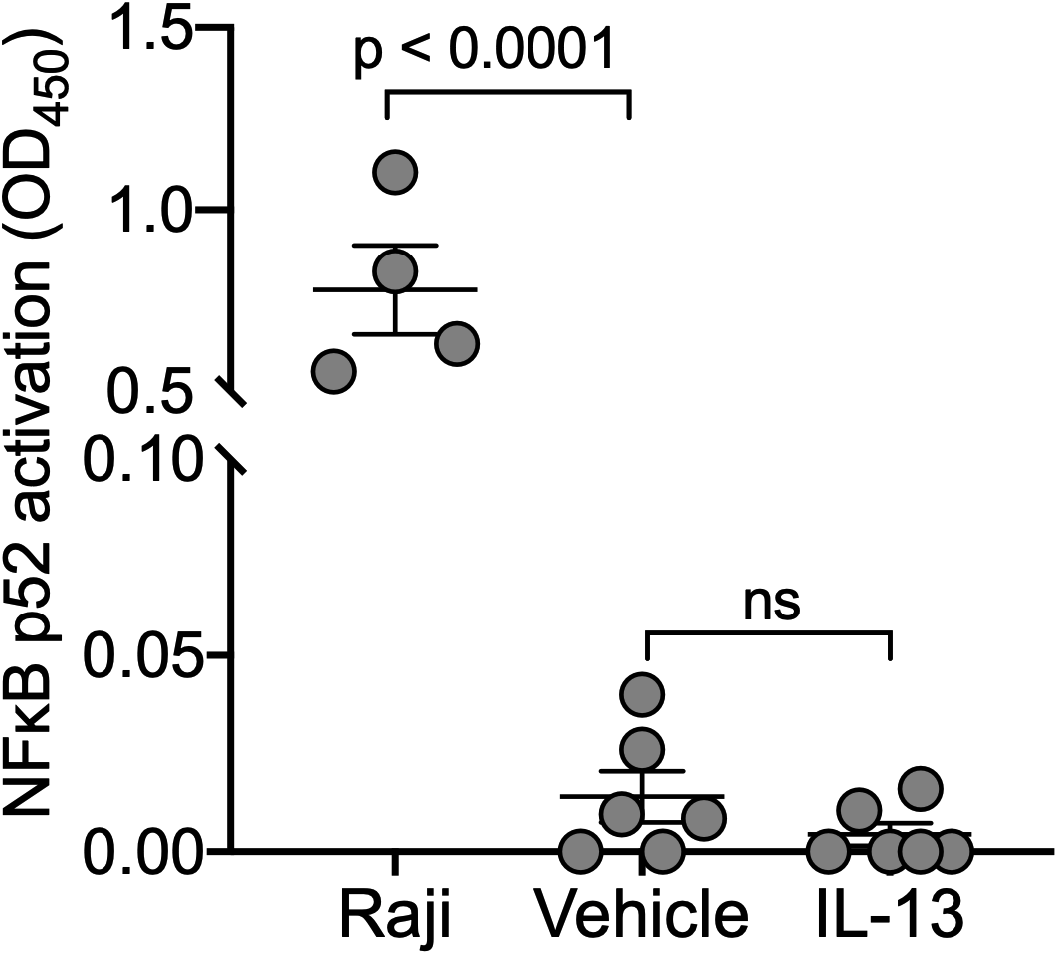
IL-13 does not induce DNA binding of p100/p52 subunit in human airway epithelial cells *in vitro*. To determine whether IL-13 induces DNA binding of the NFκB p52 subunit, primary human airway epithelial cells were treated with vehicle or IL-13 for 1 hour. Nuclear extracts were collected and IL-13-induced DNA binding of p52 subunit was measured using an ELISA-based DNA-binding assay. Each data point corresponds to epithelia from a different subject for vehicle- and IL-13-treated samples. (*n* = 6 biological replicates). Data represents the mean ± SEM, ns = not significant by paired T-test.

### IL-13 does not induce nuclear translocation of p52 in human airway epithelial cells in vitro

Previous reports have described phosphorylation of p100 S869 to be essential in p100 processing. However, we did not detect induction of p52 DNA-binding in our cultured airway epithelial cells. We suspected that the assay was not sensitive to detect p100 processing in primary airway cells. Thus, we used another method to test IL-13-induced p100 processing in human airway epithelial cells. Nuclear extracts were collected from airway epithelial cells exposed to vehicle or IL-13 for 1 hour. Then, we performed a Western blot for p52 (Fig. 5). We found that IL-13 did not induce nuclear translocation of p52 in airway epithelial cells. These findings suggest that although phosphorylation of p100 S869 is essential, it is not sufficient to induce p100 processing in airway epithelial cells. Indeed, it has been described that phosphorylation of multiple residues on p100 are required for inducible p100 processing (33).

**Figure 5.**
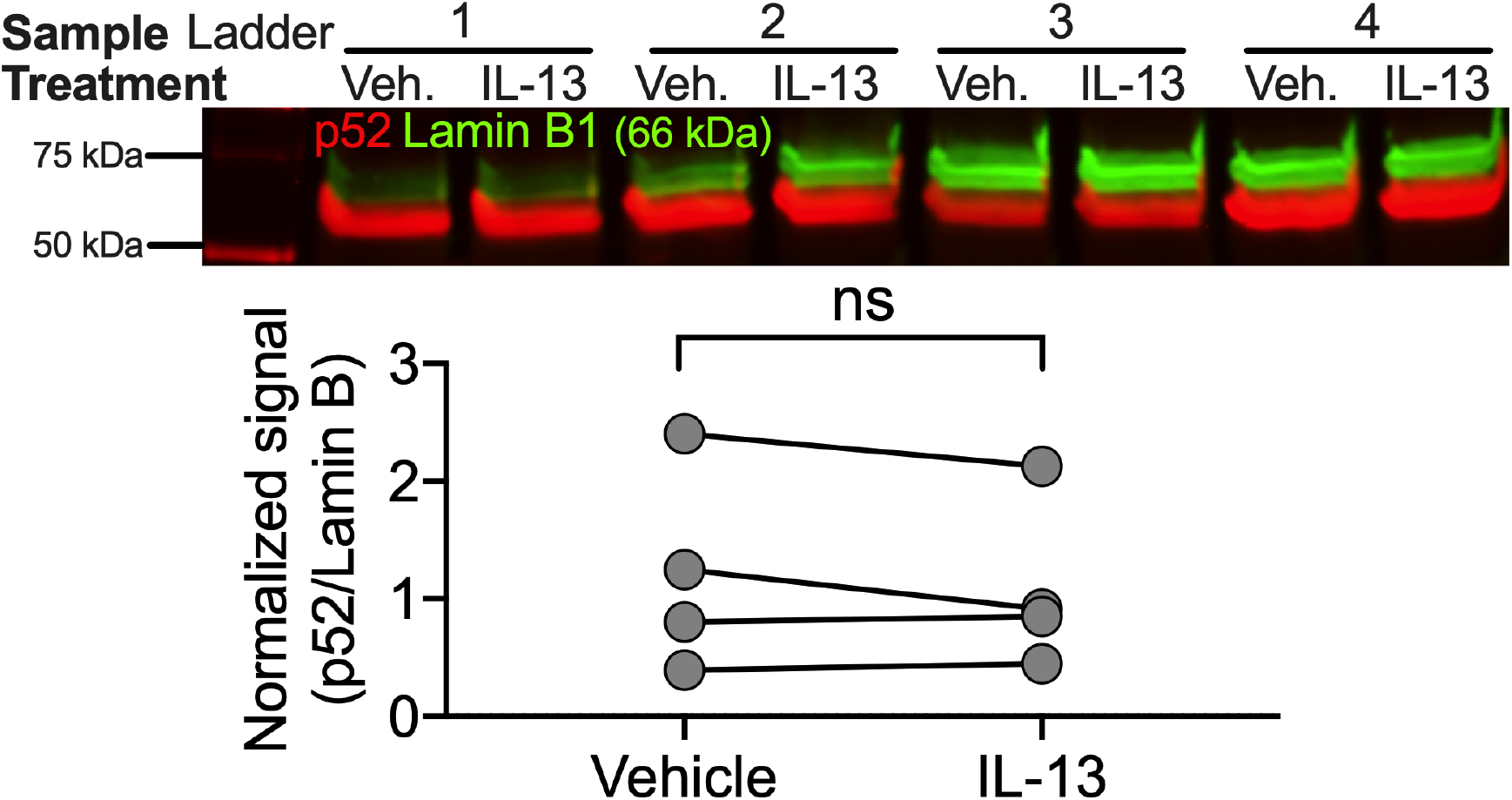
IL-13 does not induce nuclear translocation of NF*κ*B subunit p100/p52 in primary human airway epithelia *in vitro*. Primary human airway epithelial cells were treated with vehicle or IL-13 for 1 hour. To evaluate nuclear translocation of p52, nuclear proteins were isolated from the whole-cell lysate, and analyzed through Western blot. Data shown are p52 signal intensity normalized to Lamin B. n = 4 biological replicates. ns = not significant by paired T-test.

### HSP90 inhibition blocks IL-13-induced phosphorylation of p100 S869 in primary human airway epithelia in vitro

To validate our observations in our antibody array data regarding p100 phosphorylation, we tested whether IL-13-induced phosphorylation of p100 S869 can be detected through immunofluorescence staining and whether it can be blocked by geldanamycin and prevent goblet cell metaplasia. Human airway epithelial cells were treated with vehicle, HSP90i, IL-13, or IL-13 plus HSP90i for 28 days. The cells were then fixed, and immunofluorescence staining for p100 pS869 and MUC5AC (a marker for goblet cells) was performed (Fig. 6). We found that at baseline, p100 pS869 and MUC5AC were not detectable, and exposure to HSP90i alone had no effect. IL-13 treatment induced an increase in MUC5AC staining, as we expected. Consistent with our prior findings, treatment of HPS90i with IL-13 blocked this significant increase in MUC5AC (13). Moreover, exposure to IL-13 induced phosphorylation of p100 S869. However, when HSP90i was added, IL-13-induced p100 S869 phosphorylation was abolished. This provides further support that the NF*κ*B pathway is involved in airway goblet cell metaplasia induced by IL-13.

**Figure 6.**
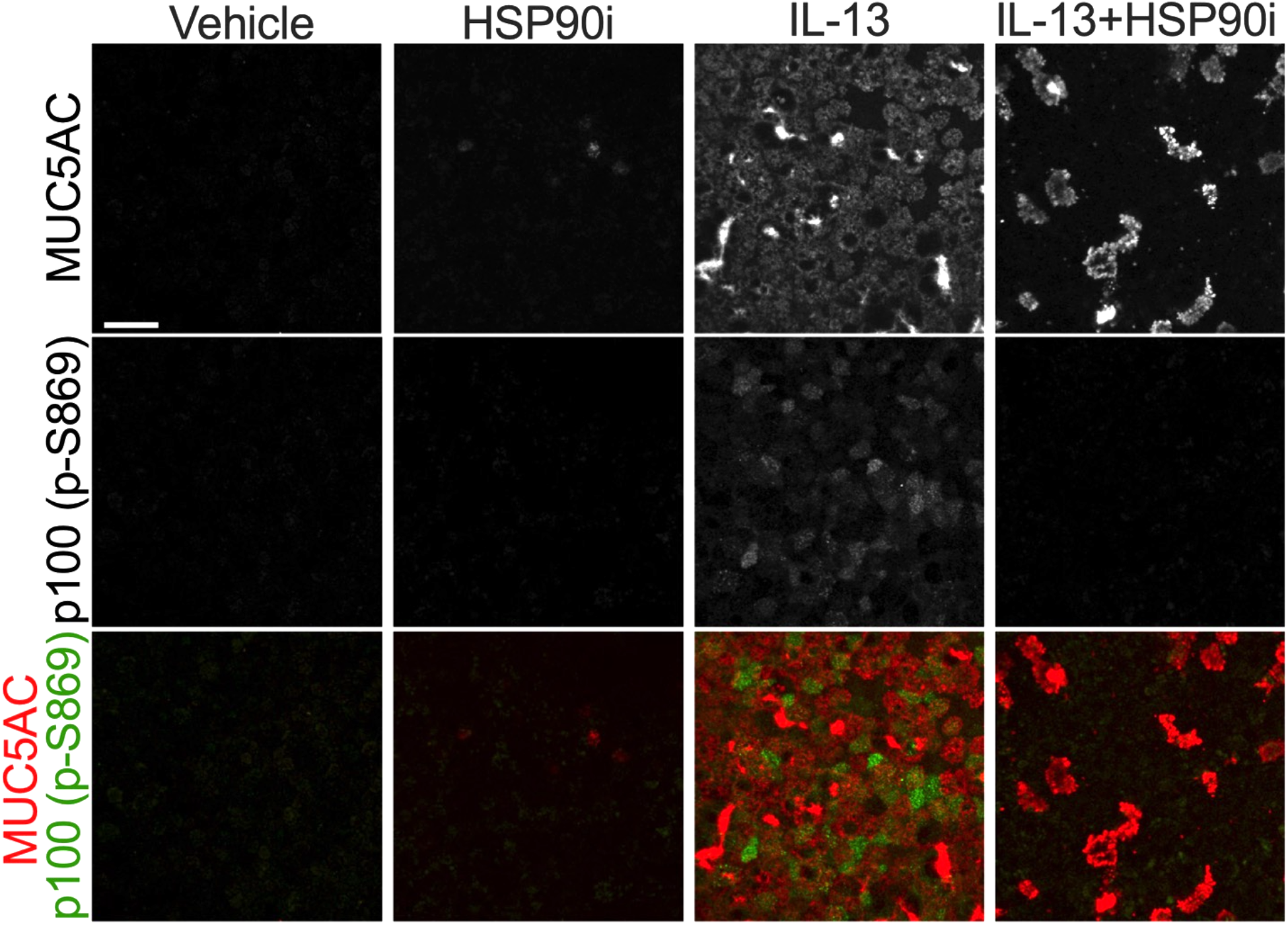
HSP90 inhibition blocks IL-13-induced phosphorylation of NF*κ*B subunit p100 in primary human airway epithelia. Primary human airway epithelial cells were treated with vehicle, HSP90 inhibitor, IL-13, or IL-13 plus HSP90 inhibitor for 28 days. Immunofluorescence staining was performed for phosphorylated Ser869 on p100 and MUC5AC. Scale bar = 30 μm.

### IKK inhibition prevents IL-13-induced airway goblet cell metaplasia

HSP90 has previously been described to play a critical role in the expression and formation of IκB kinase (IKK), an essential activator of NF*κ*B (34,35). We have demonstrated that HSP90 inhibition blocks IL-13-induced p100 S869 phosphorylation and goblet cell metaplasia. If HSP90 is required for goblet cell metaplasia and NF*κ*B activation through IKK, and NF*κ*B activation is required for goblet cell metaplasia, then NF*κ*B inhibition would block goblet cell metaplasia. Thus, we hypothesized that IKK inhibition would prevent IL-13- induced goblet cell metaplasia. To test this, we exposed airway epithelial cells to vehicle, IL-13, or IL-13 plus 10μM IKK-16, an IKK inhibitor, for 28 days. The cells were then fixed and immunofluorescence staining for MUC5AC was performed (Fig. 7A). Four representative images were acquired per condition, per donor and MUC5AC-positive cells were quantified (Fig. 7B). We found that IKK inhibition blocked IL-13-induced increase in MUC5AC-positive cells. These data strongly suggest that NF*κ*B activation is essential for the development of IL-13-induced goblet cell metaplasia in human airway epithelial cells *in vitro*.

**Figure 7.**
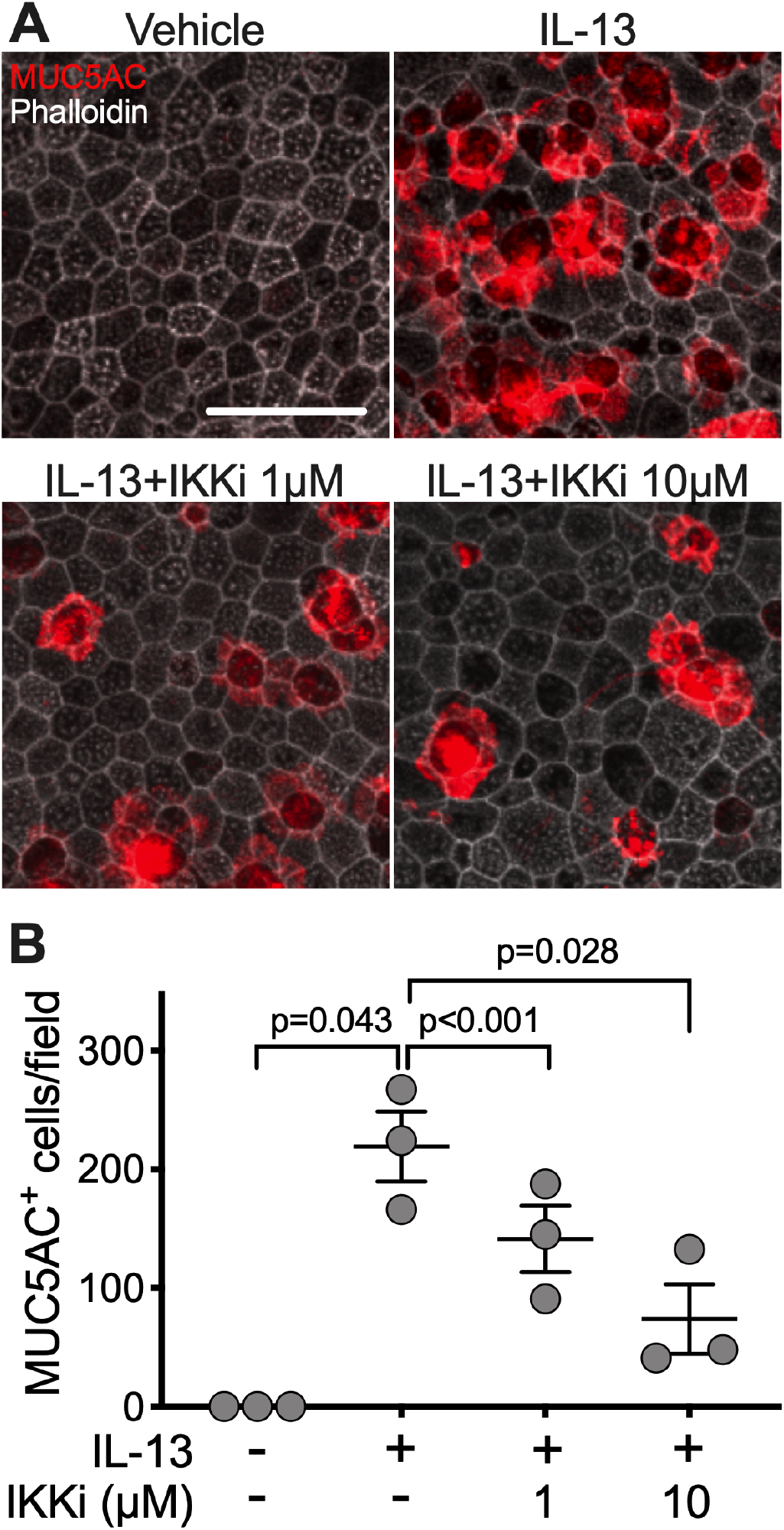
IKK inhibition prevents IL-13-induced airway goblet cell metaplasia in human airway epithelial *in vitro*. Primary human airway epithelial cells were treated with vehicle, IL-13, or IL-13 plus IKK inhibitor, IKK-16 (10uM) for 21 days. To determine whether IKK inhibition blocks IL-13-induced airway goblet cell metaplasia, immunofluorescence staining was performed for MUC5AC (goblet cells) and with phalloidin (F-actin) (**A**). Scale bar = 30 μm. MUC5AC-positive cells were quantified (**B**). Each data point corresponds to epithelia from a different subject (*n* = 3 biological replicates). Data represent the mean ± SEM. p = repeated measures one-way ANOVA with Tukey’s adjustment.

## Discussion

Our previous findings that HSP90 inhibition reverts both IL-17- and IL-13-induced goblet cell metaplasia, two distinct pathways, led us to investigate the common HSP90 client protein(s) essential for goblet cell metaplasia. Our antibody array results showed that while phosphorylation of several proteins was modulated by HSP90 inhibition in IL-13-treated airway cells, modulated phospho-sites were enriched in NFκB. Further analysis suggested a potential role of NF*κ*B in IL-13-induced airway goblet cell metaplasia. Though NF*κ*B signaling in immune cells in the context of inflammation has been well characterized, the role of NF*κ*B in the development of goblet cell metaplasia in human airway epithelia is understood less broadly (26,27).

NF*κ*B is known to affect a broad range of processes including inflammation, proliferation, and immune regulation (reviewed in (28,29)). The NF*κ*B family proteins consist of p100, p105, RelA, RelB, and cRel which form dimers and can be induced to translocate into the nucleus and affect transcription of genes. Tight regulation of this pathway involves several processes including proteasomal processing (e.g. p100, p105) and constitutive inhibition by IκB. Regulation by IκB occurs via its association with the DNA binding domain of NF*κ*B preventing nuclear translocation and binding to DNA. NF*κ*B activation via both canonical and non-canonical pathway require the action of IκB kinase (IKK), a complex consisting of catalytic subunits (IKKα and/or IKKβ) and a regulatory subunit (IKKγ also known as nuclear factor-kappa B essential modulator, NEMO) (36). The IKK complex is responsible for phosphorylation and subsequent ubiquitination and degradation of IκB, freeing NF*κ*B dimers to translocate into the nucleus and affect gene expression of various proinflammatory mediators. In the non-canonical pathway, NF*κ*B stimulation triggers activation of NF*κ*B inducing kinase (NIK) to phosphorylate and activate the IKK complex (37). The IKKα subunit is responsible for the phosphorylation and subsequent proteasomal processing of p100 into p52. In addition, NIK has also been shown to directly induce p100 processing, independent of IKK (38). In contrast, p105 processing into p50 occurs constitutively and IKK phosphorylation results in p105 degradation.

The blockade of both IL-13-induced p100 phosphorylation and goblet cell metaplasia in airway epithelial cells by HSP90 inhibition suggests a novel role of NF*κ*B in human airway epithelia. HSP90 is a molecular chaperone responsible for modulation of several signaling pathways including that of NF*κ*B (39). HSP90 modulation of the NF*κ*B signaling pathway is best demonstrated by its critical role in the stabilization of IKK, the primary regulator of NF*κ*B (34,36). Thus, the observed blockade of IL-13- induced p100 phosphorylation by HSP90 inhibition is consistent with previously described role of HSP90 in IKK function and expression (40). These data point to IKK as the potential HSP90 client required in airway goblet cell metaplasia. Indeed, IKK inhibition with IKK-16 prevented IL-13-induced goblet cell metaplasia in primary human airway epithelial cells. The prevention of IL-13-induced airway goblet cell metaplasia by inhibition of either IKK or HSP90 suggests a key regulatory step in NF*κ*B activation that require the function of both molecules. This regulatory step is, thus, also a requirement for airway goblet cell metaplasia development.

NF*κ*B activation results in the nuclear translocation of NF*κ*B dimers where it binds DNA and affect transcription. Although treatment with IL-13 induced p100 phosphorylation, we did not detect nuclear translocation or DNA binding of p52. A potential explanation for these findings may be attributed to the inhibitory activity of p100. In addition to serving as a precursor to p52, p100 can also bind other NF*κ*B proteins including its processed product acting in a similar mechanism to IκB, thereby inhibiting nuclear translocation (41). It is likely that IL-13-induced p100 phosphorylation by IKK results in proteasomal processing to p52. Unlike p100, p52 lacks the inhibitory domain and thus, the ability to bind and sequester NF*κ*B dimers in the cytosol. Therefore, IL-13-induced goblet cell metaplasia may not directly involve p100/p52 as a transcription factor, rather as a modulator of NF*κ*B activation. Our findings that IL-13 induces p100 phosphorylation but not p52 nuclear translocation or DNA binding support this mechanism. Alternatively, it is possible that phosphorylation of multiple specific residues on p100 are required for processing and IL-13-induced phosphorylation of p100 S869 is not sufficient to induce cleavage (30). Nevertheless, our data describe a novel role of IKK and HSP90 in IL-13-induced airway goblet cell metaplasia.

Despite advances in knowledge of the mechanisms of airway inflammation and epithelial remodeling, no treatments exist for airway goblet cell metaplasia. Identifying modulators of goblet cell metaplasia that can be targeted to treat chronic muco-inflammatory airway diseases has continued to be a challenge, partly due to our limited understanding of the various pathways that result in goblet cell metaplasia (42). Our findings help to understand the role of an important signaling pathway in goblet cell metaplasia induced by diverse upstream signaling mechanisms. Modulators of HSP90 and NFκB signaling are therefore potential therapeutic candidates to treat chronic muco-inflammatory diseases of the lungs.

## Experimental Procedures

All **reagents and antibodies** are listed in detail in Supporting information.

### Cell Culture and Treatment

Cells were obtained from the University of Iowa Cells and Tissue Core. Airway epithelial cells were isolated from the trachea and bronchi of human lung tissues. They were seeded onto collagen-coated, semipermeable membranes (0.6cm^2^ Millicell-PCF; MilliporeSigma), and grown at the air-liquid interface. To allow for appropriate differentiation, the epithelial cells were used at least 14 days post-seeding. A 1:1 mixture of DMEM/F12 supplemented with 2% Ultroser G (PALL Corp.) was used for the culture medium, and was changed every 3-4 days. Treatment conditions included IL-13 (20 ng/ml, R&D Systems), HSP90 inhibitor geldanamycin (25 μM, Sigma-Aldrich), and IKK inhibitor IKK-16 (10 μM, Abcam). Treatments were added to the basolateral media, of which 20 μl was transferred to the apical surface to allow basolateral and apical exposures. Treatment design and durations varied between experiments and are indicated in the results section. Reagent details are provided in the Reagents and Antibodies section.

### Phospho-Explorer Array analysis

#### Sample preparation

Protein extracts were collected from primary human airway epithelial cells. Lysate concentration was measured using Qubit Protein Assay Kit (Invitrogen) and Qubit 4 Fluorometer, then standardized across experimental groups with the target protein amount of 50 μg. Protein extraction and the antibody array was performed according to the protocol provided in the Phospho Explorer Antibody Array (Full Moon BioSystems).

#### Imaging

The microarray slides were imaged using GenePix 4000B Microarray Scanner and the signal intensities were quantified using the software GenePix Pro 6.1.

#### Data analysis

The GenePix Array List (GAL) file provided by Full Moon BioSystems was used to generate GPR files in GenePix using the recommended settings. The R/Bioconductor package “PAA” (version 3.8) (43) was used to background-correct (method “normexp”/”saddle”) and quantile-normalize the data. Further analysis was done as recommended by Full Moon BioSystems in Microsoft Excel to generate phospho-ratios. Data is available in Supporting Information files.

### Imaging Assays

#### Immunofluorescence

Cultured primary human airway epithelial cells were washed with PBS (Gibco), fixed with 4% paraformaldehyde (Electron Microscopy Sciences) in PBS for 15 minutes, permeabilized with 0.3% Triton-X (Thermo Scientific) in PBS for 20 minutes, blocked with 2% BSA (Research Products International) in SuperBlock (Thermo Scientific) for 1 hour at room temperature, incubated with the primary antibody apically overnight at 4°C, and then incubated with the secondary antibody for 1 hour at 37°C protected from light, both diluted in 2% BSA in SuperBlock. The cultures were mounted on Superfrost Plus Microscope Slides (Fisher Scientific) in Vectashield with DAPI (Vector Laboratories) and then coverslipped. The primary antibodies used were mouse anti-MUC5AC (1:5000, Invitrogen) and rabbit anti-phospho NF*κ*B p100/p52 Ser869 (1:100, Invitrogen). The secondary antibodies used were goat anti-rabbit and goat anti-mouse conjugated to Alexa Fluor 488 or 568 (1:1000, Invitrogen). To distinguish individual cells for quantification of MUC5AC-positive cells, we also used the actin stain phalloidin conjugated to Alexa Fluor 647 (1:100, Invitrogen) during the secondary antibody incubation. Reagent details, including antibody catalog and lot numbers, are provided in the Reagents and Antibodies section.

#### Imaging

All immunofluorescence studies were imaged using Olympus FluoView FV1000 confocal microscope with UPLSAPO 60X oil lens, and all confocal images were processed using the Olympus FV315-SW program.

#### Quantification of confocal images

TIFF files were formatted from Z-stacks of the original confocal images. The number of cells positive for MUC5AC was manually counted using the Cell Counter plugin within the Fiji distribution of ImageJ software. For each sample, four random images were taken per culture, one from each quadrant, and the counts were averaged.

### Western Blot

#### Nuclear lysate preparation

Nuclear extracts were isolated from cell lysates of primary human airway epithelial cultures using a technique based on a previously described cell fractionation method (44). The cells were washed with cold PBS (Gibco) twice, then first lysed with 1% NP-40 (Thermo Scientific) in PBS with phosphatase inhibitor (Sigma-Aldrich) and protease inhibitor (Sigma-Aldrich). The cells were scraped using a pipette tip, transferred to a clean microcentrifuge tube, vortexed for 10 seconds, incubated on ice for 10 minutes, then centrifuged at 14,000 × g for 30 seconds to pellet the nuclei. The nuclear pellet was washed twice by resuspending with 1% NP-40 in PBS, centrifuging, and discarding the supernatant. The nuclear pellet was resuspended with Laemmli buffer with phosphatase and protease inhibitors, sonicated at level 2 for 5 seconds, twice, and centrifuged at 14,000 × g for 5 minutes. The supernatant containing the nuclear contents was transferred to a clean tube.

#### Immunoblotting

Nuclear lysates were mixed 1:1 with 2X homemade loading buffer, boiled at 90°C for 5 minutes with mixing, and cooled in ice for 2 minutes. The samples were then subjected to SDS-PAGE at a constant voltage of 100 V for 1 hour at room temperature and transferred to nitrocellulose membrane (BIO-RAD) at a constant voltage of 30 V at 4°C overnight. The nitrocellulose membrane was blocked with 5% BSA (Research Products International) in PBS with 0.1% Tween-20 (Research Products International) for 1 hour, incubated with primary antibody for 2 hours, then incubated with secondary antibody for 1 hour. The membrane was washed three times with 1X TTBS for 10 minutes in between incubations. All incubations were done at room temperature with rocking. The primary antibodies used were rabbit anti-NF*κ*B p100/p52 (1:1000, Cell Signaling) and mouse anti-Lamin B1 (1:1000, Invitrogen). The secondary antibodies used were donkey anti-rabbit or anti-mouse conjugated to IRDye 800CW or 680LT. A Kaleidoscope protein ladder was used. Reagent details, including antibody catalog and lot numbers, are provided in the Reagents and Antibodies section.

#### Imaging and Quantification

The western blots were imaged using LI-COR Odyssey and band signal intensities were quantified using the LI-COR Image Studio Software.

### Enzyme-Linked Immunosorbent Assay (ELISA)

Primary human airway epithelial cells were treated with either vehicle or IL-13 (20 ng/ml) both apically and basolaterally for 1 hour. The cells were then lysed, and nuclear extracts were isolated using the reagents and protocol provided in the Active Motif Nuclear Extract Kit (Active Motif). IL-13 effects on NF*κ*B p52 DNA binding was evaluated using the TransAM NF*κ*B Transcription Factor Assay Kit (Active Motif). The samples were run in duplicates and absorbance was measured using a spectrophotometer at 450 nm.

## Data availability

All data generated for this manuscript is available in the results or as supporting information.

## Acknowledgments

We’d like to thank Lynda Ostedgaard (University of Iowa), Tom Moninger (University of Iowa), and Paola Vermeer (Sanford Health) for helpful discussion and/or technical support.

## Funding

This work was supported by the National Institutes of Health [NHLBI K01HL140261] (AAP); the Parker B. Francis Fellowship Program (AAP), and the Cystic Fibrosis Foundation University of Iowa Research Development Program (Bioinformatics Core) (AAP and ALT). The content is solely the responsibility of the authors and does not necessarily represent the official views of the National Institutes of Health.

## Conflict of interest

The authors declare that they have no conflicts of interest with the contents of this article.

## Footnotes

### Abbreviations (CONFIRM DEFINED)

HSP90: heat shock protein-90
HSP90i: heat shock protein-90 inhibitor
T_h_2: T helper cell type 2
IL-13: interleukin 13
IL-17: interleukin 17
NFκB: nuclear factor kappa-light-chain-enhancer of activated B cells
IKK: IκB kinase
IKB: Nuclear factor of kappa light polypeptide gene enhancer in B-cells inhibitor
Akt: Protein kinase B
Jak/STAT: Janus kinase/signal transducer and activator of transcription
IRS: insulin receptor substrate

